# Diverse genome organization strategies for polysaccharide utilization in the oceans

**DOI:** 10.64898/2026.09.21.753266

**Authors:** Aaron Oliver, Sheila Podell, Eric E. Allen

**Author notes:** Address correspondence to Eric E. Allen.

## Abstract

Carbohydrate-active enzymes (CAZymes) drive the turnover of polysaccharides in the oceans. However, the scale of diversity in polysaccharide utilization and genome organization strategies has yet to be characterized, particularly in marine ecosystems. In this paper, we introduce mpcgcdb.com as an interactive web catalog for marine CAZyme gene clusters, their associated metagenome-assembled genomes, and relevant enzyme families found in marine metagenomic samples. This database contains nearly 290,000 marine CAZyme gene clusters from more than 22,000 genomes, enabling user-initiated exploratory visualizations for genome organization networks and enzyme phylogeny. This tool will assist researchers in developing hypotheses about gene function, connecting species to metabolic niches, and identifying gaps where future work is needed to understand different pathways in marine glycan cycling.

## Introduction

The oceans contain more than 600 gigatons of dissolved organic carbon^1^ as Earth’s largest carbon sink^2^. The carbon shapes global climate^3^ as dissolved organic matter that is the core of the microbial loop^4^ powering marine microbial metabolism. Prior estimates indicate dissolved glycans make up at least 10% of dissolved organic carbon stock^5^ (with estimates north of 70% in regions of algal blooms^6^). Whatever glycans are not degraded by water column microbes sink into the deep oceans, sequestered for centuries^7^. While the biosynthesis pathways for marine polysaccharides are generally well-understood^8–10^, studies of degradative processes often consist of linking single enzymes (e.g. ^11,12^) or bacterial species (e.g. ^13,14^) to single substrates. Comprehensive understanding of marine glycan cycling requires moving beyond current paradigms of individual species-substrate relationships, heavily shaped by what microbes are culturable^15^ and what substrates are commercially available^16^, towards more comprehensive pan-oceanic analyses.

The structural complexity of individual glycans and extracellular polysaccharide matrices shape overall carbon bioavailability in the oceans^17^. Common macroalgal polysaccharides such as carrageenan, ulvan, and fucoidan are affixed with sulfate groups^18^ that must be removed by specialized enzymes before their component monosaccharides become bioavailable to the microbiome as a whole. Thus, bacteria that decompose marine polysaccharides into monomeric components often encode suites of substrate-specific carbohydrate-active enzymes (CAZymes) and sulfatases in their genomes, in some cases totaling as much as 16% of a microbe’s coding sequences^19^. Decomposition of complex macroalgal polysaccharides requires stepwise reaction pathways enabled by enzyme co-regulation, often accomplished through the colocalization of carbohydrate-active enzymes and sulfatases within the genome, forming operon-like regions known as CAZyme gene clusters (CGCs)^20–22^.

The most prolific degraders of macroalgal polysaccharides are found in the Bacteroidota, Gammaproteobacteria, Firmicutes, and the PVC (Planctomycetota, Verrucomicrobiota, Chlamydiota) superphylum^4–6^. Some species, such as *Pontiella desulfatans*^19^ or *Zobellia galactanivorans*^26^, dedicate a large proportion of their genome to degrading various marine polysaccharides, while in other instances members of a wider community each encode a complete or partial pathway dedicated to a single polysaccharide, suggesting a microbial division of labor^27^. CGCs consist primarily of CAZymes and, for sulfated substrates, sulfatase enzymes, alongside auxiliary proteins such as transporters, transcription factors, regulatory elements, and peptidases^28^. CAZymes are divided into classes by their mechanism of action and further subdivided into families based on experimentally validated activities^29^; the classes most relevant to marine polysaccharide degradation are glycoside hydrolases (GHs), which hydrolyze glycosidic bonds, and polysaccharide lyases (PLs), which cleave them non-hydrolytically^30^. Sulfatases (S1) may remove sulfate groups before, during, or after glycosidic bond cleavage, and initial desulfation is required before CAZymes can bind some highly sulfated polysaccharides^31^. Carbohydrate esterases (CEs) similarly remove acetyl and methyl groups to improve accessibility for other enzymes^32^.

Polysaccharide utilization loci were first described in 2006 to characterize the genomic architecture of *Bacteroides thetaiotaomicron* strains that degraded human milk oligosaccharides^33^. In the Bacteroidota, the sugar transporter SusC, the glycan-binding protein SusD, and their homologs^34^ are hallmark genes that allow rapid computational identification of putative polysaccharide degradation gene clusters. As such, the first computational annotation tools and databases were restricted to the Bacteroidota phylum^35–38^.

The first automated tool for prediction of polysaccharide utilization loci was developed in 2015 as PULDB^36^, using a small set of terrestrial Bacteroidota genomes primarily associated with humans. The first software program able to annotate loci in user-provided genomes was PULpy, released in 2018^37^. That same year, sulfatase annotations were added to a new version of PULDB^38^, enabling identification of CGCs targeting sulfated polysaccharides; however, this update still focused solely on the Bacteroidota. Advances in annotation robustness subsequently revealed many gene clusters lack the SusC/SusD markers once thought necessary for polysaccharide degradation, and gene-cluster identification is now centered on colocalized CAZymes within CAZyme gene clusters^35^.

The most recent attempt to catalog marine CGCs, dbCAN-seq^22,39^, is built from the CGC-Finder module of dbCAN^28,40^. It is the first database to include both marine-specific genomes and genomes outside the Bacteroidota, enabling investigations of whole-microbiome function. Its substrate-prediction module subdivides CAZymes into substrate-specific activities, but the performance of this feature lacks validation, especially for marine organisms; for example, the marine-specific section of dbCAN-seq contains more CGCs predicted to degrade human milk oligosaccharides (62) than the marine polysaccharides alginate (47), agarose (32), or carrageenan (3).

At the same time large collections^41–43^ of diverse genomes from marine environments have become increasingly available, but not yet applied to perform a pan-oceanic analysis of marine carbon metabolism. The most comprehensive set of marine genomes in a single database to date is the Global Ocean Microbiome Catalog^44^ (GOMC), which contains more than 43,000 genomes across more than 100 prokaryotic phyla. There is rising interest in leveraging these expansive genome databases to understand marine metabolism^46^ and discover new enzymes of industrial relevance^47^.

Our prior work has shown that many of the enzymes required to degrade algal polysaccharides are not encoded in terrestrial ecosystems^48^ and that the diversity of CAZymes and sulfatases encoded in marine metagenomes are largely uncaptured by current databases^24^. Thus, although incorporating sulfatase annotations into identification of CGCs is especially crucial to understand the diversity of marine polysaccharide degradation strategies, it is not yet addressed by currently available CGC prediction tools. In this manuscript, we introduce mpCGCdb (mpcgcdb.com), an interactive web catalog that integrates 289,962 marine CAZyme gene clusters from 22,607 dereplicated metagenome-assembled genomes. These clusters are organized by taxonomic lineage and encoded enzyme families, to enable user-directed exploration of polysaccharide-utilization and genome-organization strategies across the global marine microbiome and aid in the discovery of novel enzymes, metabolic niches for bacterial families, and functions for uncharacterized auxiliary genes.

## Methods

### Genome selection

24,195 dereplicated genomes were downloaded from the Global Ocean Microbiome Catalog^44^ (GOMC; https://db.cngb.org/maya/datasets/MDB0000002), along with their predicted proteins. These genomes were high quality as determined by CheckM^49^ v1.0.12 (median 87.2% completeness, 0.99% contamination) and this genome set incorporates previous pan-oceanic metagenome surveys such as *Tara* Oceans^50^. Genomes were assigned taxonomic lineages using the Genome Taxonomy Database Toolkit^51^ v. 2.1.1, based on the Genome Taxonomy Database^42^ r207.

### Cluster annotation

To annotate tens of thousands of metagenomic genomes at scale, we developed mpCGC (Marine Polysaccharide CAZyme Gene Clusterer), distributed as a Nextflow^52^ pipeline (github.com/AaronAOliver/mpCGC). To identify, mine, and summarize large numbers of CAZyme gene clusters predicted to target marine polysaccharides, the software incorporates three data flows (**Figure 1**).

**Figure 1.**
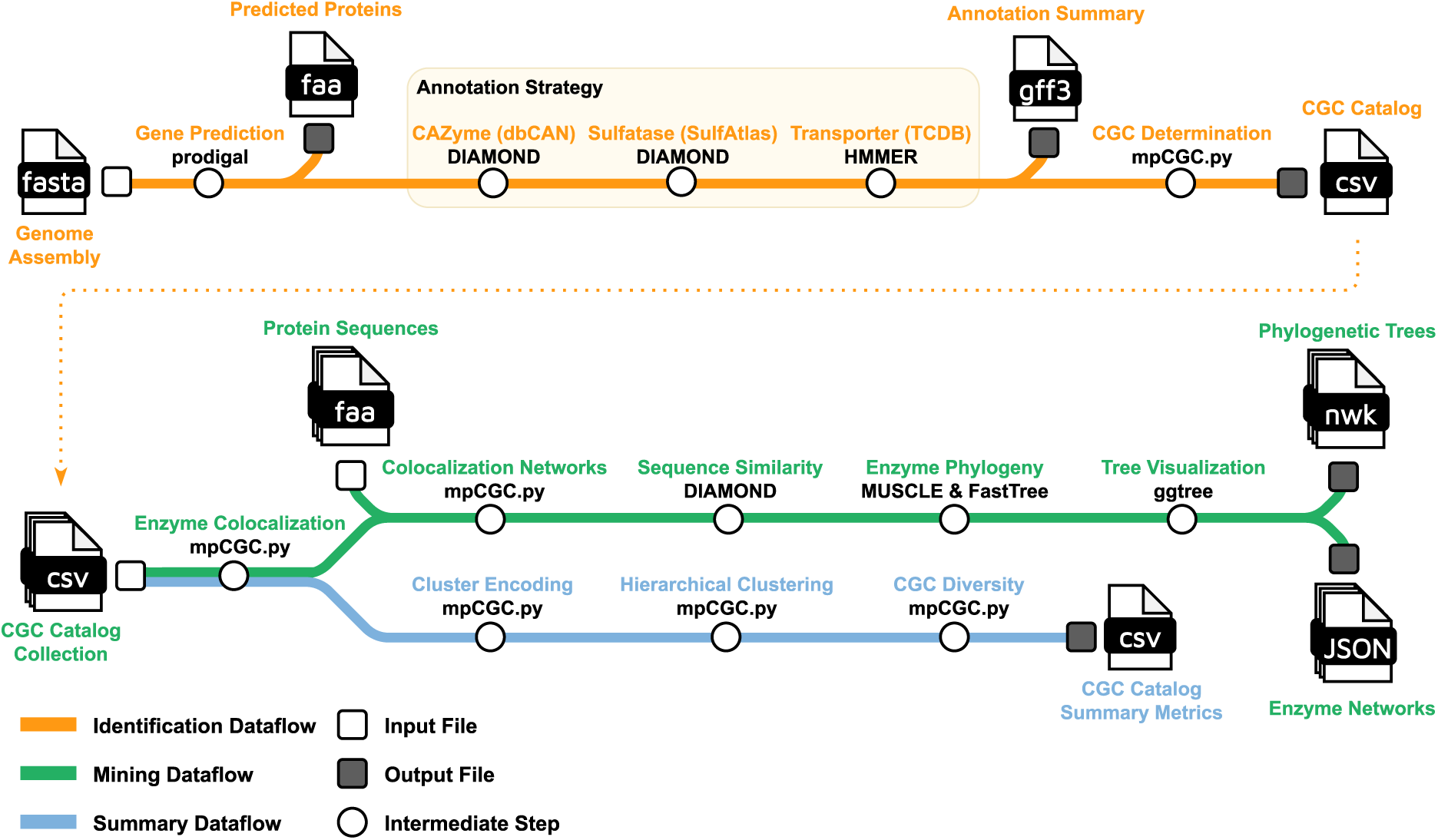
mpCGC pipeline workflow. mpCGC is composed of three dataflows: annotation of CAZyme gene clusters (orange), mining collections of CGCs for network and phylogenetic visualizations (green) and generating summary and diversity statistics for annotated CGCs (blue). The mpCGC pipeline can either begin from fasta files, genomic/metagenomic contigs or catalogs of CGCs previously generated from mpCGC or CGC-Finder.

The initial identification data flow functions similarly to tools such as dbCAN’s CGC-Finder^28,40^ and PUL prediction methods^37,38^. Given a query genome or genome set, proteins are predicted with Pyrodigal^53,54^ v.3.7.1. A triple-annotation strategy then assigns (i) carbohydrate-active enzyme families from the dbCAN database^40^ v. 13 via DIAMOND^55^ BLASTP v. 2.1.9 and PyHMMER^56,57^ v. 0.12.0, (ii) sulfatase families from the SulfAtlas database^58^ v. 2.3.1 via DIAMOND BLASTP, and (iii) transporters from the Transporter Classification Database (TCDB)^59^ for ABC transporters and the SusC/SusD complex. A seed-and-extend algorithm builds clusters from co-localized CAZymes and transporters, extending the initial gene cluster in both directions using a user-provided gene-boundary extension limit, following prior algorithmic approaches.

### Network and phylogenetic analyses

The mining dataflow of mpCGC searches collections of CAZyme gene clusters for patterns in genome organization and signals associated with taxonomic lineage to create an annotated sequence similarity network. An all-vs-all pairwise DIAMOND BLASTP search is performed using the parameters -k 0 -e 1e-30 to generate a sequence similarity network^60^ for all genes in the CGC catalog using e-value to generate weighted edges for pairwise distance. An e-value of 1e-30 was chosen^61^ as the cutoff for edge formation. The genes in the sequence similarity network are clustered into communities using the Leiden community identification algorithm^62^ with resolution = 1. For network communities in the master sequence similarity network that were not annotated by mpCGC, the most central member by degree centrality was annotated using InterProScan^63^. This sequence similarity network and associated node classifications are then subset across either taxonomic lineages or the presence of specific marker genes to provide colocalization networks. For example, the master sequence similarity network can be subset to use only genes from Verrucomicrobiota CGCs (https://mpcgcdb.com/taxonomy/dbacteria/pverrucomicrobiota/) or CGCs that contain at least one GH2_7 β-galactosidase (https://mpcgcdb.com/GH2_7/). Community subsets of the sequence similarity network form nodes in sequence colocalization networks as previously described in https://github.com/AaronAOliver/KiritimatiallesSCoNe. For each catalytic CAZyme and sulfatase subfamily, we have pre-calculated sequence similarity networks using the subset of all genes colocalized with that subfamily, as well as all CGC genes from each taxonomic phylum, class, and order.

Annotated CAZymes and sulfatases for a user’s input are automatically built into phylogenetic trees using the alignment tool MUSCLE^64^ v. 5.1 and tree building software FastTree^65^ v. 2.1.11. Publication-quality visualizations are generated for all phylogenetic trees using R v. 4.5.2 packages ggtree^66,67^ and ggtreeextra^68^. An additional command-line option enables the generation of custom phylogenetic trees with user-selected network communities or external gene lists as input.

The summary dataflow branch of mpCGC estimates diversity of predicted CAZyme gene clusters using a hierarchical clustering of pairwise hamming distance^69^ between clusters. CGCs are encoded using a binary presence-absence of CAZyme and SulfAtlas annotations (S1, GH, PL, and CE). Two clusters are considered overlapping in function if one cluster contains a superset of all annotations of the other cluster. Diversity metrics can be calculated using random subsets of genomes or CGCs to recreate rarefaction-style analyses to account for size of the original genomic or metagenomic input contigs.

## Results

mpCGCdb (mpcgc.com) has become the newest and most comprehensive database of marine CAZyme gene clusters, containing more than 15x more genomes and CGCs than the previous state-of-the-art dbCAN-seq marine^22^ (**Table 1**). The addition of sulfatase annotations allows mpCGCdb to resolve clusters targeting sulfated marine polysaccharides, such as carrageenan, fucoidan, and ulvan, that are systematically missed by CAZyme-only pipelines. Both carbohydrate and sulfate components of the mpCGC database dwarf the size of earlier Bacteroidota-specific PULDB^38^ and PULpy^37^ databases that were limited by requiring SusC/SusD marker genes for cluster annotation. The large number of phyla included in our database, present at sufficient genome counts to make robust scientific inferences, enable comparative marine glycogenomics not feasible using data from prior studies. We will discuss the construction and organization of the web database, followed by a set of vignettes showing how this data can be used to generate novel biological hypotheses.

**Table 1.** Comparison of existing CAZyme gene cluster and polysaccharide utilization loci databases.

|  | Oliver<br>*et al.* 2026 | Zheng<br>*et al.* 2023 |  | Terrapon<br>*et al.* 2018 | Stewart<br>*et al.* 2018 |
| --- | --- | --- | --- | --- | --- |
| Database Name | mpCGCdb | dbCAN-seq |  | PULDB | PULpy |
| Included Genomes | Marine:<br>22,607 | Total:<br>9,421 | Marine:<br>1,496 | 2,494 | 5,414 |
| Predicted Clusters | 289,962 | 168,906 | 18,367 | 81,235 | 96,117 |
| CAZyme Aware | Yes | Yes |  | Yes | Yes |
| Sulfatase Aware | Yes | No |  | Yes | No |
| Predict Substrates | No | Yes |  | No | No |
| Taxon Agnostic | Yes | Yes |  | No | No |
| Public Software | Yes | Yes |  | No | Yes |
| CGC Download | Yes | Yes |  | No | Yes |

### Database and web design

To create the curated database, we used 22,607 dereplicated metagenome-assembled genomes with at least 1 CGC, with taxonomy spanning 139 phyla, 337 classes, 988 orders, 2,044 families, and 4,579 genera with the marine polysaccharide CAZyme gene clusterer (mpCGC) pipeline (**Figure 1**). This analysis identified 289,962 CGCs that contain 406 glycoside hydrolase families, 98 polysaccharide lyase families, 18 carbohydrate esterase (CE) families, and 102 sulfatase subfamilies. **Table 2** summarizes the distribution of genomes, coding sequences, and CAZyme gene clusters from the most abundant bacterial phyla in the database, defined as having more than 300 genomes each.

**Table 2.** Database statistics for the most abundant bacterial phyla. Data columns represent the number of genomes associated with each phylum, the number of predicted proteins associated with each phylum, the number of predicted CAZyme gene clusters, the percentage of proteins associated with a CAZyme gene cluster compared to all proteins, and the total numbers of CAZymes and sulfatases predicted for phylum-associated CAZyme gene clusters.

| Phylum | MAGs | CDS | Total CGCs | CGCs (% Genome) | CAZymes in CGCs | Sulfatases in CGCs |
| --- | --- | --- | --- | --- | --- | --- |
| Proteobacteria | 9,349 | 23,332,809 | 113,137 | 3.49% | 155,232 | 13,923 |
| Bacteroidota | 3,304 | 8,260,007 | 64,705 | 5.57% | 107,080 | 14,013 |
| Actinobacteriota | 1,011 | 3,164,895 | 19,989 | 2.98% | 28,652 | 2,302 |
| Planctomycetota | 993 | 3,619,198 | 14,436 | 2.04% | 18,478 | 4,187 |
| Chloroflexota | 955 | 2,187,494 | 7,220 | 1.02% | 9,002 | 536 |
| Cyanobacteria | 722 | 1,771,514 | 13,244 | 6.22% | 17,083 | 810 |
| Patescibacteria | 595 | 495,329 | 1,263 | 1.31% | 1,737 | 44 |
| Verrucomicrobiota | 589 | 1,639,386 | 7,300 | 2.84% | 9,212 | 2,672 |
| Desulfobacterota | 516 | 1,439,982 | 5,020 | 2.13% | 6,814 | 217 |
| Marinisomatota | 452 | 704,953 | 2,298 | 1.98% | 3,293 | 302 |
| Campylobacterota | 419 | 682,829 | 2,606 | 2.17% | 3,579 | 126 |
| Firmicutes | 309 | 1,067,317 | 9,242 | 7.50% | 14,280 | 1,101 |
| Acidobacteriota | 306 | 918,543 | 3,491 | 2.38% | 4,476 | 486 |

mpCGCdb is split into multiple views that subdivide the available data and visualizations. As shown in **Supplemental Figure 1**, the navigation sidebar on the home page previews distinct views of the database, dividing CAZyme gene clusters by enzyme family, taxonomic lineage, source genome, and individual clusters. Each enzyme family page provides interactive phylogeny tools to study enzyme evolution and identify potential horizontal gene transfers, a sequence similarity network and sequence-colocalization network to connect auxiliary genes to enzyme substrate and function, and taxonomic distributions to aid in future isolation and characterization. **Supplemental Figure 1a** shows an example sequence similarity network download page and sequence colocalization network for the β-agarase/β-porphyranase class GH86. Taxonomic lineage pages provide an overview of CGC content, a sequence similarity network, a sequence colocalization network, a ranked table of co-occurring subfamilies, and lineage-specific downloads. **Supplemental Figure 1b** shows the sunburst plot for interactive navigation of the mpCGCdb website by taxonomic lineage.

**Supplemental Figure 1. Page layout of mpCGCdb.** The homepage navigation sidebar (left) with webpage screenshots of (A) enzyme-specific, (B) taxonomy-specific, (C) genome-specific, and (D) gene cluster-specific views of the marine polysaccharide CAZyme gene cluster database.

For each annotated genome, we provide the following metadata: Genome Taxonomy Database^42^ lineage, CheckM^49^ completeness and contamination, genome size, GC content, and coding density. Each page also contains a downloadable table of every CGC in the genome, and direct links to each gene cluster’s detail page. **Supplemental Figure 1c** shows an example page and table of annotated CGCs for the spirochaete *Winmispira thermophila* (GCA_000184345.2), originally isolated from a marine hot spring and known to degrade alginate, xylan, and laminarin^70^. An example gene arrow diagram is shown in **Supplemental Figure 1d**, for CGC #4 from *Winmispira thermophila*.

The downloads page (https://mpcgcdb.com/downloads/) contains bulk downloads for all CGC, CAZyme, and sulfatase annotations for the entire dataset, hosted on Zenodo (record 20219287; 10.5281/zenodo.20219284). The global sequence similarity network, network communities, and similarity co-localization network are provided as tab-separated node and edge lists, with full Cytoscape sessions on Zenodo (record 20214548; 10.5281/zenodo.20214547). Amino acid FASTA files for individual enzyme families, taxonomic lineages, and individual CGCs are available on their respective web pages. High level statistics about the entire mpCGC database are provided at https://mpcgcdb.com/statistics. The most abundant CAZyme families inside annotated CGCs are the α-amylase family GH13, the large multifunctional GH3 family, and the lytic transglycosylase GH23. The most abundant sulfatase subfamilies were S1_6, S1_8, and S1_2. More than 3,000 genomes contained more than 25 CGCs, composing 12.5% of the original Global Ocean Microbiome Catalog.

### Mining biological insights from mpCGCdb

This large collection of CAZyme gene clusters from more than 100 phyla enables comparisons of degradative potential between marine lineages that have not previously been possible. One example is a quantification of CGC richness between the most abundant phyla in mpCGCdb. **Figure 2** shows that Firmicutes, Bacteroidota, and Planctomycetota contain the richest collection of CGCs in our dataset, while Cyanobacteria, Desulfobacterota, and Archaea show the least diversity in genome organization strategies for CAZymes. Using Chao2 richness estimation, we predict more than 9,000 unique CAZyme and sulfatase arrangements in marine Bacteroidota, more than 4,000 in Planctomycetota, and fewer than 400 in Cyanobacteria (**Supplemental Table 1**). By these estimates, fewer than 38% of marine Bacteroidota CGCs and 36% of marine Planctomycetota CGCs are captured by current metagenomic sequencing efforts.

**Figure 2.**
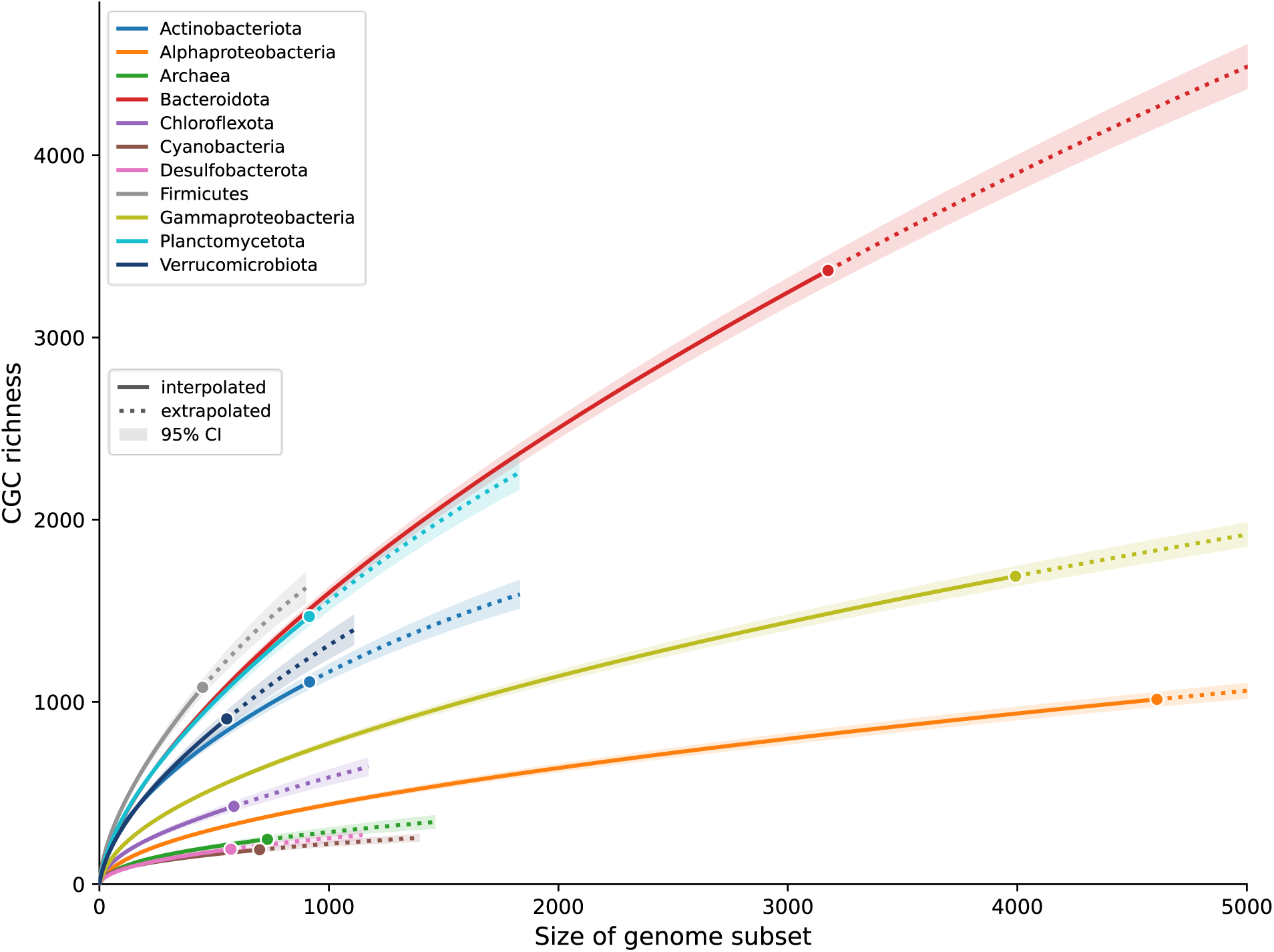
Rarefaction and extrapolation of CGC diversity across marine microbial lineages. A) Rarefaction and extrapolation curves of CGC richness as a function of the number of genomes sampled, computed independently for each taxonomic group following the iNEXT framework^71,72^. Each genome is treated as a sampling unit and each CGC is encoded using presence-absence of CAZyme and SulfAtlas families followed by hierarchical clustering. Solid lines show interpolation up to each group’s observed genome count (filled circle), and dotted lines show extrapolation. Shaded bands are 95% confidence intervals from 100 bootstrap replicates; line color denotes taxonomic group.

**Supplemental Table 1. Chao2 CGC richness asymptote estimations for marine lineages.** Data columns represent the number of genomes associated with each lineage, the number of unique CGC families, the Chao2 estimated asymptote for total CGC families, the asymptote standard error and confidence intervals, and the percentage of predicted asymptotic CGC family count recovered in mpCGCdb.

The genes encoded in these CGCs display phylum-specific signals (**Figure 3a**). Phylum membership significantly structured CGC enzyme class composition (PERMANOVA^73,74^ on Hellinger distances, pseudo-F = 169.8, R² = 0.12, p < 0.001), explaining ∼12% of compositional variation, though groups overlapped substantially rather than forming discrete clusters (mean silhouette^75,76^ ≈ 0.03). Established genomes with exceptionally high percentages of the genome related to CGCs can be found in all of the major phyla (**Figure 3b**). Selected genomes showcasing this adaptation to a diet rich in complex carbohydrates are shown in **Supplemental Table 2**. These include species noted in prior literature for their glycan degradation capabilities, such as the agar and fucoidan degrading *Saccharophagus degradans*^77^ (18.6% CGCs, Gammaproteobacteria), the thermophilic fermenter *Pyrococcus furiosus*^78^ (7.57%, Archaea), and the xylan degrading *Paenibacillus sacheonensis*^79^ (23.59%, Firmicutes).

**Figure 3.**
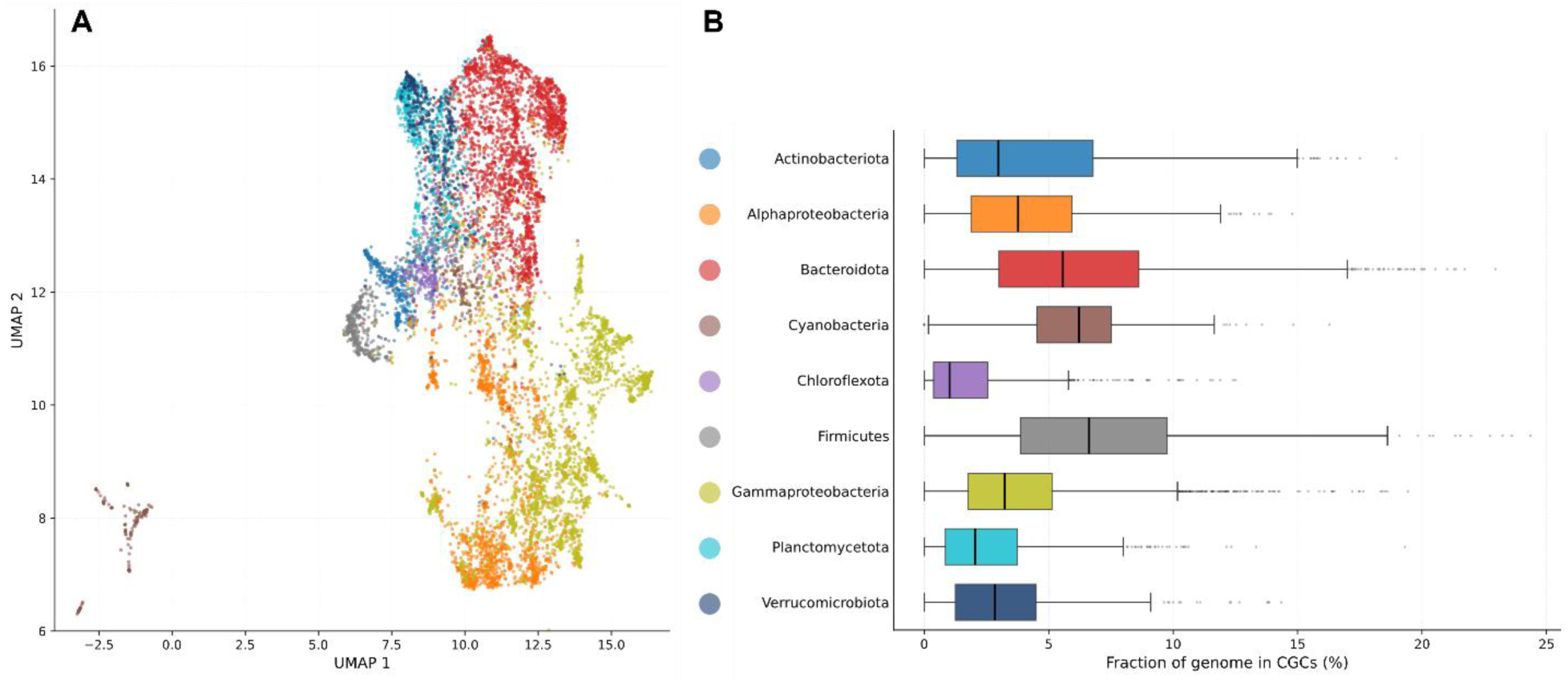
Phylum-specific signatures of carbohydrate utilization strategies in marine metagenomes. A) A UMAP^83^ dimensionality reduction of CAZyme and sulfatase genes inside annotated CGCs for genomes in mpCGCdb. B) Relative abundance of CGCs per genome, as calculated by CGC genes ÷ total genes, using the same color scheme as panel A.

This analysis simultaneously identified genomes with expanded glycan breakdown specialization whose CAZyme and sulfatase repertoires have not been previously described in literature. These include *Cohnella sp.* CIP 111063 (GCA_002237815.1, 24.3%, Firmicutes), *Mangrovibacterium lignilyticum*^80^ (GCA_010882175.1, 22.9%, Bacteroidota), *Sedimentisphaera salicampi* (GCA_002239625.1, 19.3%, Planctomycetota), *Microbacterium sp.* AISO3^81^ (GCA_002204305.1, 19.0%, Actinobacteriota), and *Tichowtungia aerotolerans*^82^ (GCA_009905215.1, 13.8%, Verrucomicrobiota). In total, 1,070 genomes in mpCGCdb (4.7% of genomes) dedicate more than 10% of their genomes to CGCs. These previously uncharacterized outlier strains offer promising targets for culturing and for the identification of novel enzymes and pathways for degrading marine polysaccharides.

**Supplemental Table 2. High quality genomes in mpCGCdb abundant in CAZyme gene cluster genes.** Data columns represent the taxonomy lineage, genome ID from the Global Ocean Microbiome Catalog, CheckM completeness and contamination, the number of CGCs identified by mpCGC and the relative abundance of CGCs as calculated by CGC genes ÷ total genes.

**Figure 4** showcases visualizations generated for the large sulfatase subfamily S1_4, an endo-D-galactose sulfate hydrolase known to act on glycosaminoglycans^84^. Subfamily S1_4 is distributed across most phyla (**Figure 4b**), with 27.8% of Firmicutes and 22% of Bacteroidota genomes containing at least one S1_4 in a CGC. However, only 6.1% of Proteobacteria genomes contain this sulfatase (**Figure 4c**). The Flavobacteriaceae family of Bacteroidota forms a distinct clade in the bottom left of the phylogenetic tree (**Figure 4a**), while other clades have wider taxonomic diversity. S1_4 colocalizes most often with sulfatase subfamilies S1_7, S1_16, and S1_8 (**Figure 4d**), but is colocalized with 120 gene classifications in more than 10 CGCs. Interactive versions of these figures are available at https://mpcgcdb.com/S1_4. These network visualizations help users generate hypotheses about associated transporter function, gene regulation through transcription factors, and enzyme function for poorly characterized or promiscuous enzyme classes.

**Figure 4.**
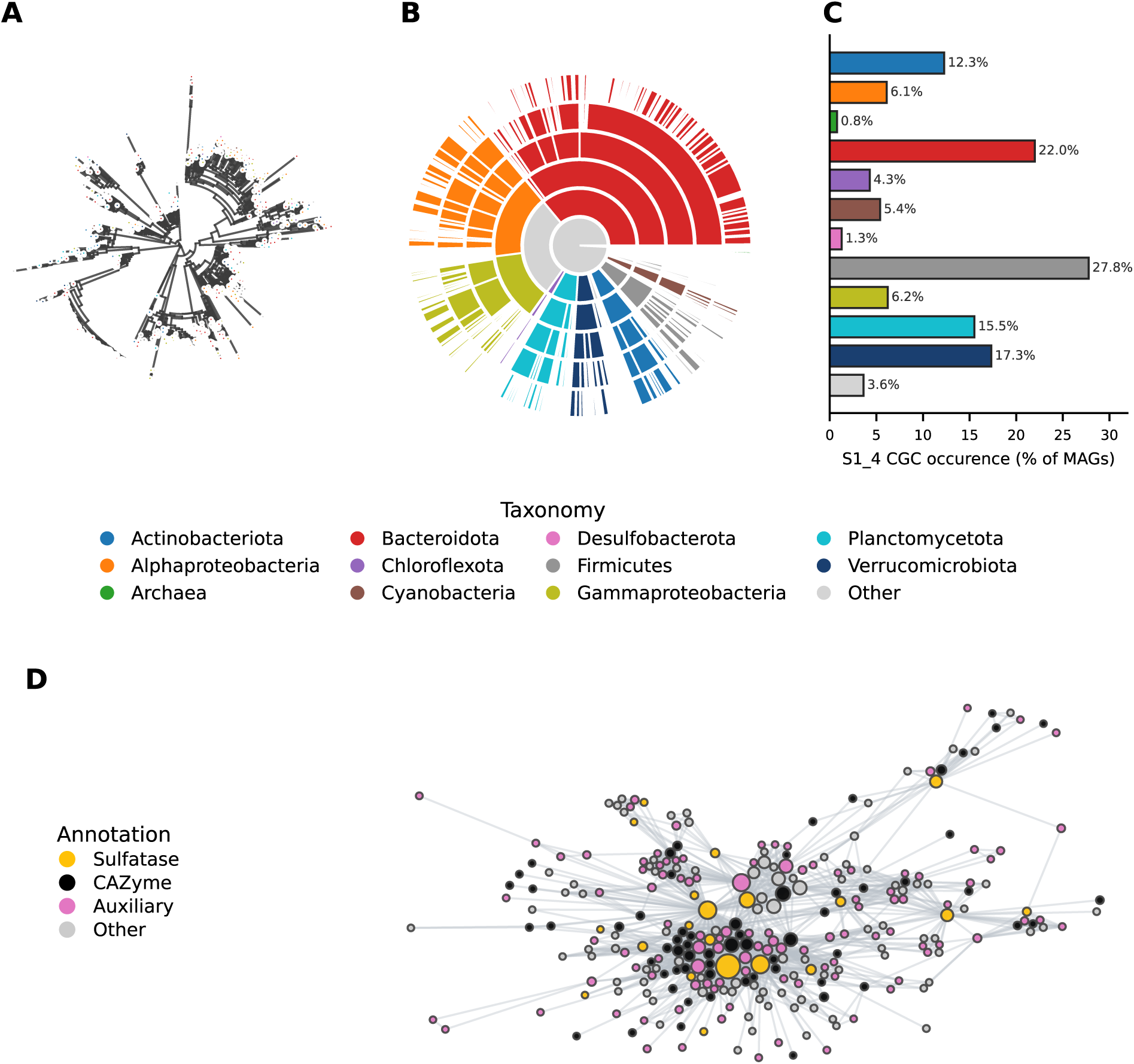
Exploring sulfatase subfamily S1_4 through interactive visualization. A) Phylogenetic tree for S1_4 sulfatase enzymes found in mpCGCdb, colored by MAG taxonomy. B) Taxonomic sunburst plot showing species-level resolution of S1_4 distribution. C) Occurrence of S1_4 in annotated CGCs across selected taxonomic groups. D) Sequence colocalization network of genes colocalized with S1_4 sulfatases using 1e-80 and 6 minimum cooccurrence cutoffs.

The collection of 289,962 CAZyme gene clusters enables exploration of enzyme classes that may be coregulated or target related substrates in degradation pathways at the largest scale described in literature. **Figure 5a** shows a colocalization frequency heatmap for glycoside hydrolase, polysaccharide lyase, and sulfatase families, highlighting enzyme classes that appear in the same CGCs. Commonly colocalized enzymes do share predicted substrate preferences, with regions of α-amylase/α-glucosidase (**Figure 5b**, **Figure 5c**), β-glucanase (**Figure 5d**), alginate and glycosaminoglycan lyase (**Figure 5e**), mannosidase (**Figure 5f**), pectinase (**Figure 5g**), arabinofuranosidase (**Figure 5h**), xylanase (**Figure 5i**), and fucosidase (**Figure 5j**) activity. Bacteroidota are particularly adapted to the degradation of α- and β-glucans compared to the other phyla (**Supplemental Figure 2**), while Verrucomicrobiota and Planctomycetota specialize in the degradation of sulfated fucans and Firmicutes genomes are relatively enriched in pectinases. Cyanobacteria CAZymes are primarily related to glycogen remodeling, with minimal utilization of any more complex sugars.

**Figure 5.**
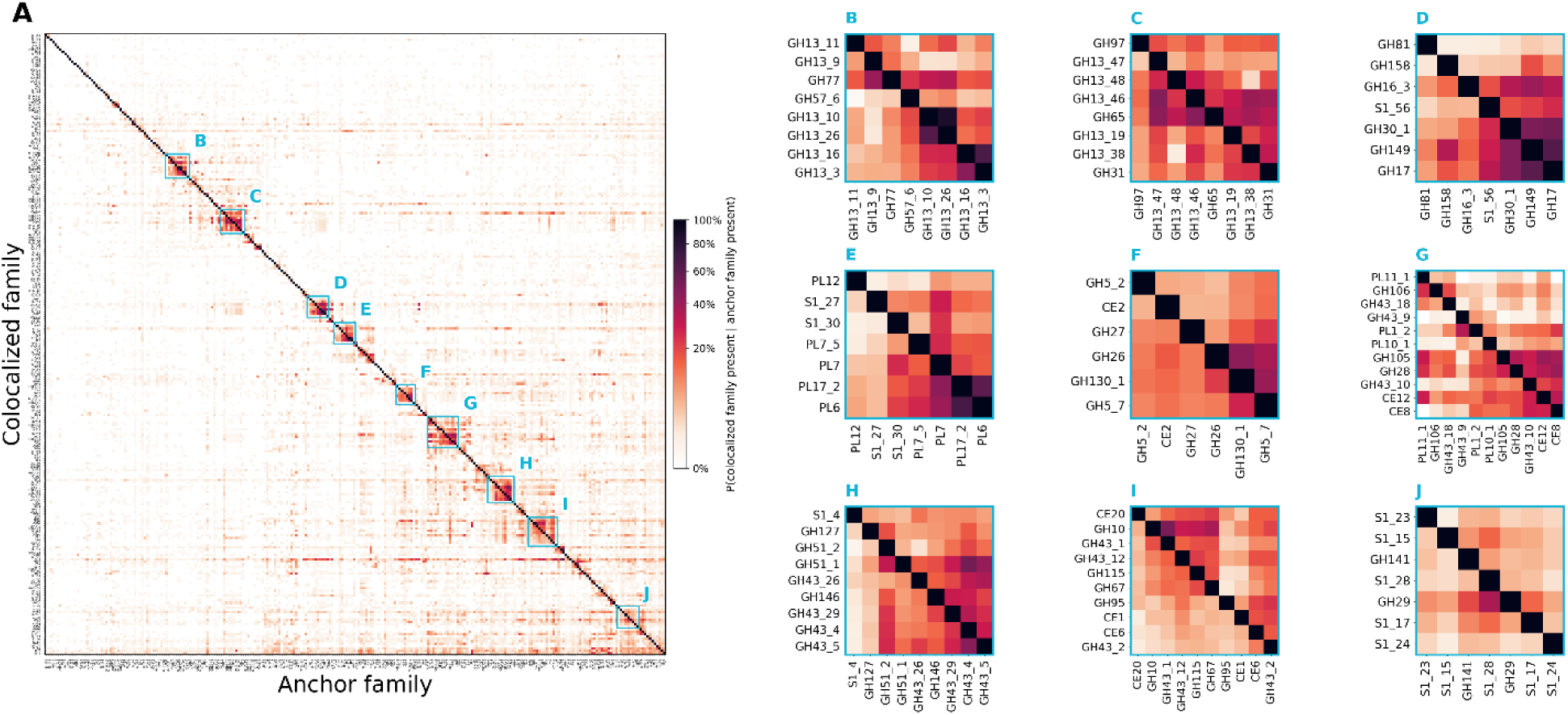
Colocalization matrix for catalytic CAZyme and sulfatase families. A) Heatmap displaying colocalization frequency for enzyme families present in at least 100 CGCs. Each cell represents the frequency that the enzyme on the y-axis is present in a CGC, given the presence of the enzyme on the x-axis (anchor). B-J) Selected regions with high colocalization frequency along the matrix diagonal, showing strong associations between groups of enzyme classes.

**Supplemental Figure 2. Taxonomic distribution of selected spatially conserved enzyme modules.** Beeswarm charts of enzyme abundance, normalized for genome count and genome size, for modules defined in the panels of Figure 5.

These consistent colocalization trends also include sulfatases with no Enzyme Commission classification in SulfAtlas, S1_56, S1_30, S1_23, S1_28, and S1_24. S1_23, S1_28, and S1_24 likely target sulfated fucans (**Figure 5j**), but S1_56 and S1_30 are most associated with enzymes predicted to act on laminarin and alginate, unsulfated brown algae polysaccharides. S1_30 is primarily found in Bacteroidota, while the other undescribed sulfatase classes have a wider taxonomic distribution (**Supplemental Figure 3**), so these enzymes may share a core role across phyla. Connecting understudied enzyme classes to taxonomic distributions can speed up isolation, characterization, and comparative genomics efforts.

**Supplemental Figure 3. Taxonomic distribution of selected sulfatase subfamilies without experimentally validated Enzyme Commission numbers.** Bar charts of abundant sulfatase subfamilies in mpCGCdb lacking EC numbers but showing strong patterns of colocalization with other enzyme classes. Numbers below enzyme classes display total enzyme count.

## Discussion

Here, we introduce mpcgcdb.com to the glycobiology and marine genomics communities as a comprehensive source of taxonomic-, genome-, enzyme- and gene cluster-centric views of marine carbon cycling through glycan degradation. Despite the vast amount of polysaccharide utilization research involving the Bacteroidota, new annotations from mpCGC show that the Actinobacteriota, Firmicutes, and PVC superphylum genomes contain similarly high diversity of CGCs. This finding suggests that novel polysaccharide utilization strategies might be found in more targeted searches for novel genomes from these phyla at a similar rate that novelty has been found and previously described in the Bacteroidota. Similarly, while only 1.2% of mpCGCdb CAZymes have no functional prediction, 23.8% of sulfatases in mpCGCdb are annotated as an enzyme class with no Enzyme Commission number. Beyond enzymes directly breaking individual glycan bonds, mpCGC has identified thousands of poorly characterized auxiliary gene families consistently colocalized within CGCs that may be key in connecting gene clusters to substrates.

It is likely that our predictions of CGC total diversity and the strength of interactions between enzyme classes are underestimates due to several factors. Fragmented assemblies from metagenomic sequencing may disperse genes in the same CGC across multiple contigs, limiting the number of long, complex CGCs that appear in MAGs. A focus on classified hydrolase and sulfatase enzymes overshadow other key genes, for example oligosaccharide transporter diversity may overemphasize pathway substrate specificity^85^. The field of glycobiology’s prior focus on terrestrial polysaccharides has overshadowed the diversity of marine CGCs as a source of novel enzymes and genome organization strategies^48^, and taxonomic bias in recovering genomes from environmental sequencing^86^ may further contribute to lower annotation sensitivity for rare taxa and CGCs targeting less abundant substrates. Nonetheless, these predictions provide the first baseline for understanding the richness of microbial glycan cycling strategies in the ocean.

## Conclusion

mpCGC and mpCGCdb provide a versatile new CGC detection pipeline and the most comprehensive database of marine CAZyme gene clusters assembled to date, spanning 289,962 clusters from 22,607 dereplicated genomes across 139 phyla. By incorporating sulfatase annotations and removing the Bacteroidota-specific marker-gene requirement of earlier tools, mpCGC captures polysaccharide-degradation machinery in lineages omitted by previous databases. These analyses reveal that high CGC diversity and genomes heavily invested in carbohydrate catabolism extend across the Actinobacteriota, Firmicutes, and PVC superphylum rather than being confined to Bacteroidota. A collection of interactive phylogenies, sequence-similarity networks, and similarity-colocalization networks, organized by both taxonomic lineage and enzyme family, can help researchers identify candidate enzymes, propose functions for poorly characterized sulfatases and auxiliary genes from their genomic context, and prioritize outlier strains for cultivation and biochemical validation. As marine genome catalogs continue to expand, the taxon-agnostic and reproducible design of mpCGC positions this resource to grow alongside them, so mpCGCdb may accelerate the discovery of the novel enzymes and metabolic strategies that underpin the turnover of marine polysaccharides in the global carbon cycle.

## Acknowledgements

Computational analyses were performed using the San Diego Supercomputer Center’s Triton Shared Computing Cluster. This work was funded by the United States Government, National Science Foundation grants OCE-2414798 and EF-2025217 to E.E.A., National Institutes of Health NIEHS grant P01ES035541-01 to E.E.A., and United States Department of Agriculture National Institute of Food and Agriculture Predoctoral Fellowship 2026-67011-46317 to A.O.

## Data Availability

The mpCGCdb web resource is freely available at https://mpcgcdb.com. Bulk CGC, CAZyme, and sulfatase annotations are archived on Zenodo (record 20219287). Global sequence similarity networks, network community assignments, and similarity colocalization networks (Cytoscape sessions and node/edge lists) are archived on Zenodo (record 20214548). The source metagenome-assembled genomes were obtained from the Global Ocean Microbiome Catalog (https://db.cngb.org/maya/datasets/MDB0000002). The mpCGC analysis pipeline is distributed as a Nextflow workflow at github.com/AaronAOliver/mpCGC.

