## Supplementary figures and images for "Diverse genome organization strategies for polysaccharide utilization in the oceans"

### Supplemental Figure 1

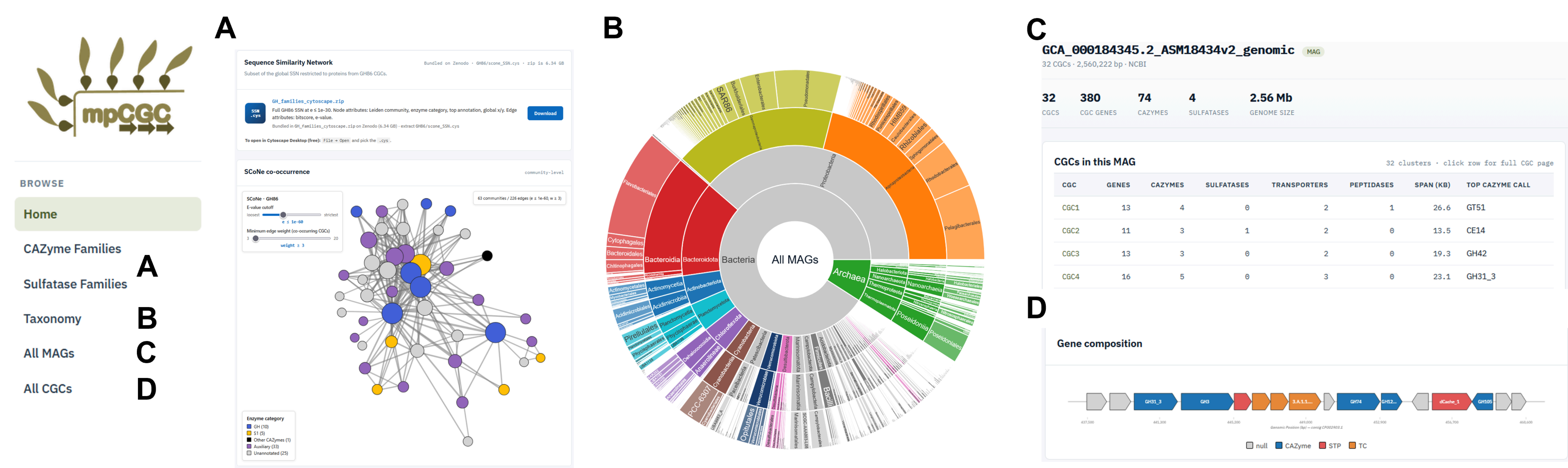

### Supplemental Figure 2

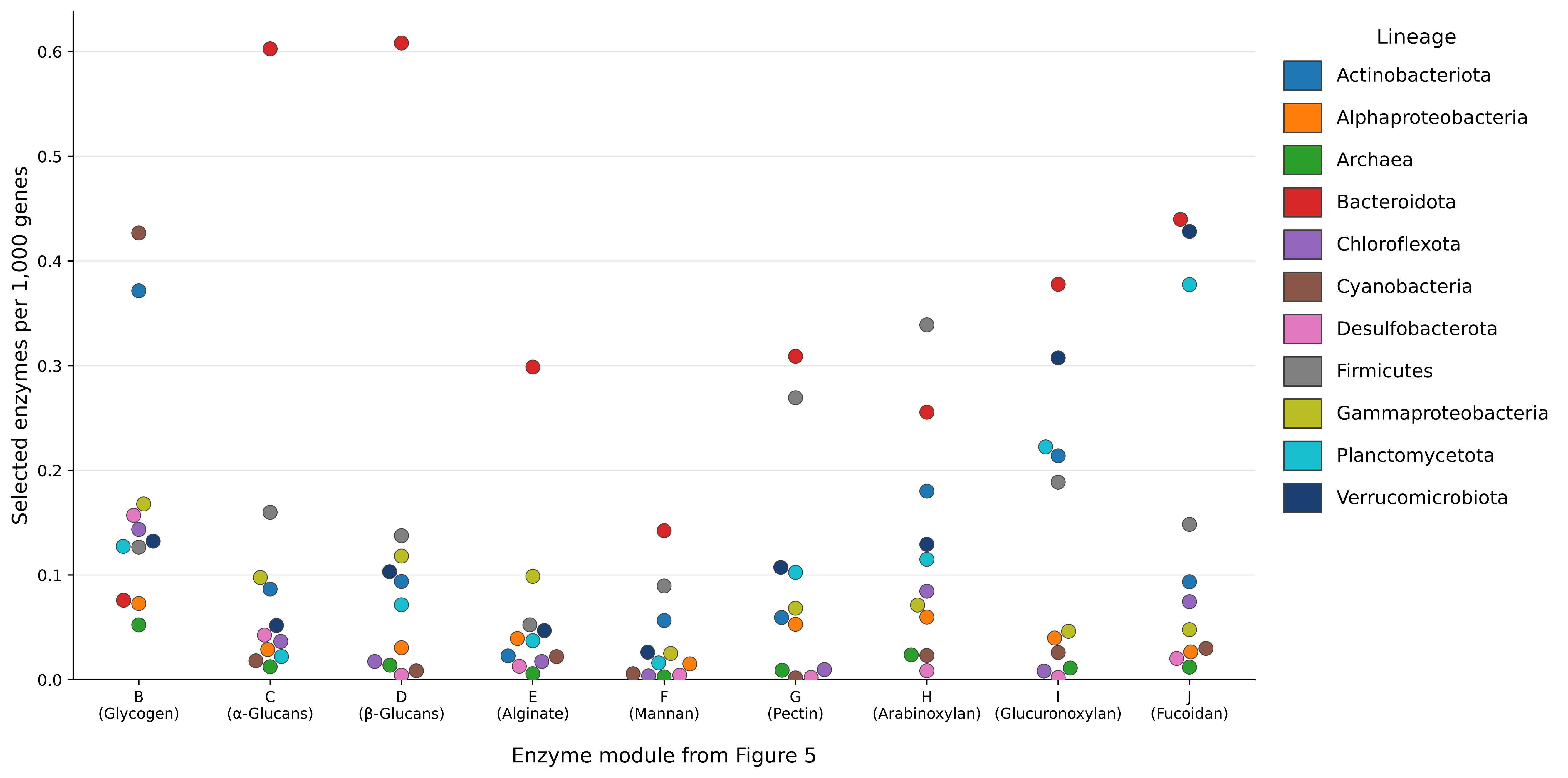

### Supplemental Figure 3

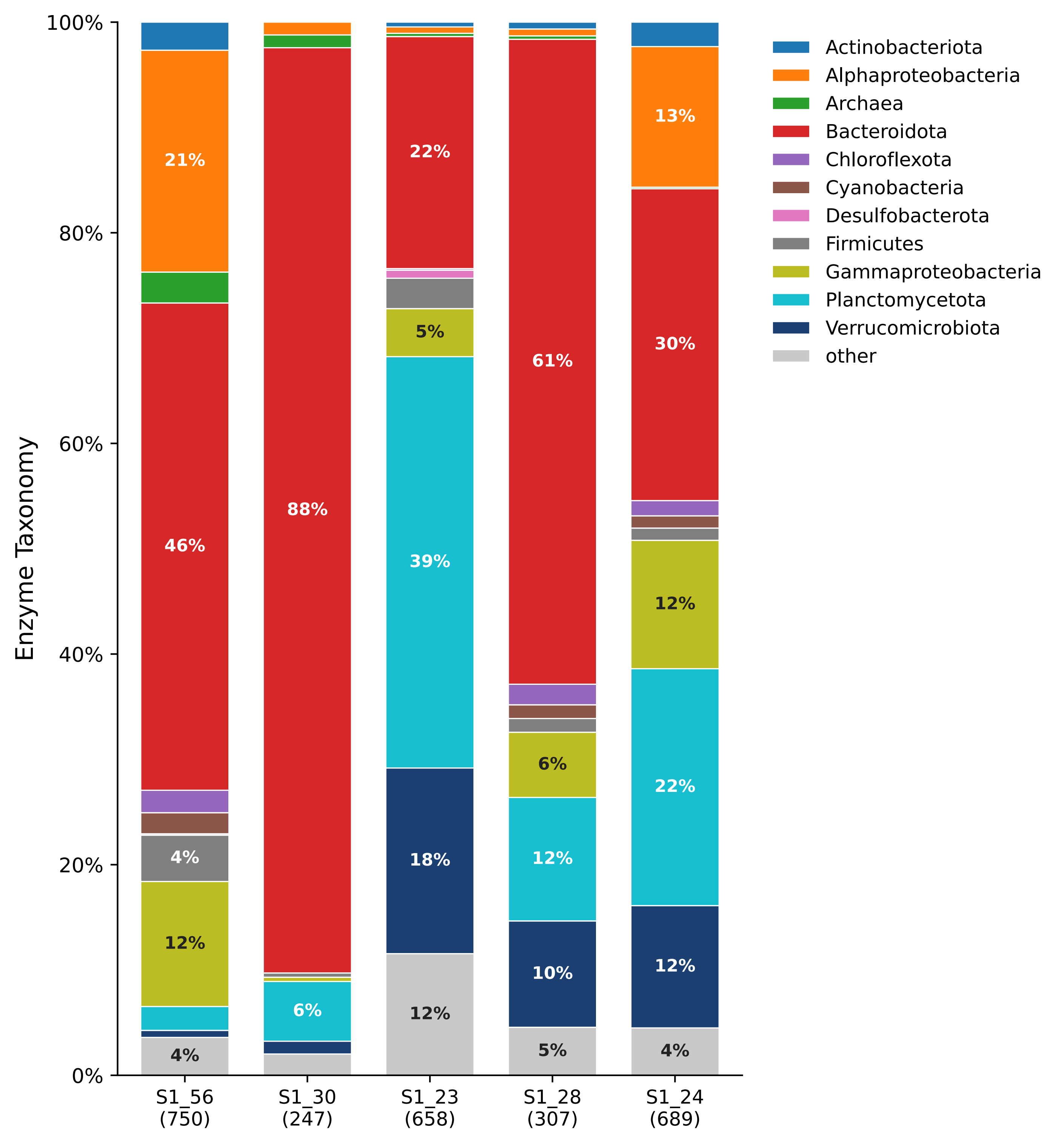
