## Supplemental Tables 1, 2 for "Diverse genome organization strategies for polysaccharide utilization in the oceans"

**Supplemental Table 1. Chao2 CGC richness asymptote estimations for marine lineages.** Data columns represent the number of genomes associated with each lineage, the number of unique CGC families, the Chao2 estimated asymptote for total CGC families, the asymptote standard error and confidence intervals, and the percentage of predicted asymptotic CGC family count recovered in mpCGCdb.

| Lineage | Genomes | Unique CGC | Asymptote | SE | 95% CI | % obs in mpCGCdb |
|---|---|---|---|---|---|---|
| **Bacteroidota** | 3,175 | 3,368 | 9,164 | 389 | 8,450–9,978 | 37% |
| **Planctomycetota** | 915 | 1,469 | 4,116 | 273 | 3,632–4,708 | 36% |
| **Gammaproteobacteria** | 3,992 | 1,690 | 3,786 | 202 | 3,426–4,219 | 45% |
| **Verrucomicrobiota** | 555 | 907 | 2,669 | 238 | 2,260–3,201 | 34% |
| **Firmicutes** | 450 | 1,080 | 2,664 | 185 | 2,340–3,070 | 41% |
| **Actinobacteriota** | 916 | 1,110 | 2,345 | 146 | 2,090–2,667 | 47% |
| **Alphaproteobacteria** | 4,608 | 1,014 | 2,294 | 161 | 2,016–2,650 | 44% |
| **Chloroflexota** | 586 | 427 | 1,350 | 201 | 1,032–1,835 | 32% |
| **Archaea** | 732 | 246 | 519 | 76 | 406–714 | 47% |
| **Desulfobacterota** | 573 | 192 | 407 | 66 | 311–578 | 47% |
| **Cyanobacteria** | 698 | 189 | 366 | 58 | 284–519 | 52% |

**Supplemental Table 2. High quality genomes in mpCGCdb abundant in CAZyme gene cluster genes.** Data columns represent the taxonomy lineage, genome ID from the Global Ocean Microbiome Catalog, CheckM completeness and contamination, the number of CGCs identified by mpCGC and the relative abundance of CGCs as calculated by CGC genes ÷ total genes.

| Lineage | Genome | GTDB Genus and Species | Comp. (%) | Contam. (%) | # CGCs | CGCs % Genome |
|---|---|---|---|---|---|---|
| Actinobacteriota | GCA_002090155.1 | g__Demequina;s__Demequina sp002090155 | 97.11 | 1.16 | 45 | 17.51 |
| Actinobacteriota | GCA_004217235.1 | g__Plantactinospora;s__Plantactinospora sp004217235 | 99.47 | 0.65 | 114 | 16.94 |
| Actinobacteriota | GCA_002090175.1 | g__Demequina;s__Demequina sp002090175 | 97.69 | 0.58 | 39 | 16.6 |
| Actinobacteriota | GCA_003856975.1 | g__Plantibacter;s__Plantibacter cousiniae | 99.49 | 0.67 | 62 | 16.57 |
| Actinobacteriota | GCA_003688855.1 | g__Arachnia;s__Arachnia antarctica | 98.52 | 0.52 | 52 | 16.33 |
| Alphaproteobacteria | GOMC.bin.15577 | g__Sphingomonas;s__Sphingomonas paucimobilis | 99.66 | 0.11 | 47 | 14.79 |
| Alphaproteobacteria | GCA_006363825.1 | g__Oceanicola;s__Oceanicola pacificus_A | 99.92 | 0.1 | 52 | 13.93 |
| Alphaproteobacteria | GOMC.bin.9689 | g__Sphingomonas;s__Sphingomonas sp002292295 | 99.66 | 0.34 | 49 | 13.85 |
| Alphaproteobacteria | GCA_001469845.1 | g__Sphingomonas;s__Sphingomonas sp000282895 | 99.48 | 0.37 | 52 | 13.45 |
| Alphaproteobacteria | GCA_009758415.1 | g__Devosia;s__Devosia marina | 99.59 | 0.68 | 44 | 13.25 |
| Archaea | GCA_002214485.1 | g__Thermococcus;s__Thermococcus pacificus | 100 | 0 | 18 | 8.89 |
| Archaea | GCA_002214365.1 | g__Thermococcus;s__Thermococcus celer | 100 | 0 | 17 | 7.88 |
| Archaea | GCA_008245085.1 | g__Pyrococcus;s__Pyrococcus furiosus | 99.5 | 0 | 17 | 7.57 |
| Archaea | GCA_002214505.1 | g__Thermococcus;s__Thermococcus siculi | 99.5 | 0 | 16 | 7.52 |
| Archaea | GCA_001317345.1 | g__Thermococcus_A;s__Thermococcus_A sp001317345 | 98.51 | 0.5 | 15 | 7.43 |
| Bacteroidota | GCA_010882175.1 | g__Mangrovibacterium;s__ Mangrovibacterium sp010882175 | 100 | 2.42 | 87 | 22.94 |
| Bacteroidota | GCA_002210225.1 | g__Geofilum;s__Geofilum rhodophaeum | 100 | 1.09 | 65 | 21.7 |
| Bacteroidota | GOMC.bin.11812 | g__Paludibacter;s__ | 93.62 | 2.69 | 62 | 21.35 |
| Bacteroidota | MARD_ SAMN05444343_ REFG_ MMP05444343 | g__Mangrovibacterium;s__ Mangrovibacterium diazotrophicum | 100 | 2.42 | 86 | 21.28 |
| Bacteroidota | GCA_000626635.1 | g__Draconibacterium;s__Draconibacterium orientale | 99.89 | 3.06 | 78 | 21 |
| Chloroflexota | GCA_003569045.1 | g__Aggregatilinea;s__Aggregatilinea lenta | 95.45 | 2.91 | 60 | 12.5 |
| Chloroflexota | GOMC.bin.7929 | g__UBA6055;s__UBA6055 sp011523545 | 90.91 | 0 | 45 | 11.59 |
| Chloroflexota | GOMC.bin.12248 | g__UBA6055;s__UBA6055 sp002238485 | 91.82 | 0 | 30 | 10.4 |
| Chloroflexota | GOMC.bin.7586 | g__JAFGMM01;s__ | 91.09 | 3.03 | 51 | 10.32 |
| Chloroflexota | GOMC.bin.9321 | g__JAAUSY01;s__ | 95.91 | 4.43 | 40 | 10.23 |
| Cyanobacteria | GCA_000316665.1 | g__Rivularia;s__Rivularia sp000316665 | 99.78 | 1.22 | 81 | 12.15 |
| Cyanobacteria | GCA_000952155.1 | g__Aliterella;s__Aliterella atlantica | 94.22 | 0.67 | 60 | 11.32 |
| Cyanobacteria | GOMC.bin.11732 | g__Pleurocapsa;s__ | 96.65 | 0.98 | 52 | 11.22 |
| Cyanobacteria | GCA_003215655.1 | g__AG-409-J16;s__AG-409-J16 sp003215655 | 98.91 | 0 | 24 | 11.07 |
| Cyanobacteria | GCA_008807075.1 | g__RSCCF101;s__RSCCF101 sp008807075 | 98.91 | 1.63 | 31 | 10.44 |
| Desulfobacterota | GCA_003402415.1 | g__Cupidesulfovibrio;s__Cupidesulfovibrio sp000226255 | 99.41 | 1.18 | 35 | 11.24 |
| Desulfobacterota | GCA_003966735.1 | g__Desulfovibrio_Q;s__Desulfovibrio_Q ferrophilus | 99.41 | 0 | 30 | 10.48 |

**Supplemental Table 2** (continued).

| Lineage | Genome | GTDB Genus and Species | Comp. (%) | Contam. (%) | # CGCs | CGCs % Genome |
|---|---|---|---|---|---|---|
| Desulfobacterota | GCA_000425245.1 | g__Maridesulfovibrio;s__Maridesulfovibrio hydrothermalis | 100 | 0 | 36 | 10.38 |
| Desulfobacterota | GCA_008369015.1 | g__Oryzomonas;s__Oryzomonas japonica | 99.35 | 0 | 35 | 10.23 |
| Desulfobacterota | MARD_ SAMN05660337_ REFG_ MMP05660337 | g__Maridesulfovibrio;s__Maridesulfovibrio ferrireducens | 100 | 0 | 35 | 9.98 |
| Firmicutes | GCA_002237815.1 | g__Cohnella;s__Cohnella sp003001675 | 99.68 | 0.22 | 146 | 24.35 |
| Firmicutes | GCA_009909195.1 | g__Paenibacillus_Z;s__Paenibacillus_Z sacheonensis | 99.67 | 2.33 | 135 | 23.59 |
| Firmicutes | GCA_009649995.1 | g__Paenibacillus_O;s__Paenibacillus_O pasadenensis | 99.54 | 0.48 | 106 | 23.23 |
| Firmicutes | GCA_009882985.1 | g__Paenibacillus_E;s__Paenibacillus_E silvestris | 99.19 | 1.06 | 133 | 22.71 |
| Firmicutes | GCA_001885765.1 | g__Enterococcus_I;s__Enterococcus_I aquimarinus | 98.51 | 0.02 | 48 | 21.96 |
| Gammaproteobacteria | GOMC.bin.5763 | g__Gilvimarinus;s__ | 99.56 | 0.24 | 69 | 19.43 |
| Gammaproteobacteria | OceanDNA-b36690 | g__Saccharophagus;s__Saccharophagus degradans | 97.41 | 0.43 | 75 | 18.61 |
| Gammaproteobacteria | OceanDNA-b35148 | g__Enterobacter;s__Enterobacter hormaechei_A | 98.21 | 0.33 | 73 | 18.49 |
| Gammaproteobacteria | TARA_ SAMEA4397586_ METAG_ ADKHDKBK | g__Gilvimarinus;s__ | 98.28 | 0.03 | 62 | 18.4 |
| Gammaproteobacteria | GCA_000463525.1 | g__Marinimicrobium;s__Marinimicrobium koreense | 99.57 | 0.48 | 57 | 18.33 |
| Planctomycetota | GCA_002239625.1 | g__Sedimentisphaera;s__Sedimentisphaera salicampi | 98.86 | 1.14 | 49 | 19.31 |
| Planctomycetota | GCA_001999965.1 | g__Limihaloglobus;s__Limihaloglobus sulfuriphilus | 97.73 | 1.14 | 43 | 13.33 |
| Planctomycetota | GCA_007859955.1 | g__Botrimarina;s__Botrimarina colliarenosi | 96.55 | 0 | 53 | 12.14 |
| Planctomycetota | GCA_007753265.1 | g__Botrimarina;s__Botrimarina mediterranea | 96.48 | 1.15 | 50 | 10.61 |
| Planctomycetota | GCA_007747445.1 | g__Poriferisphaera;s__Poriferisphaera corsica | 94.32 | 0 | 45 | 10.5 |
| Verrucomicrobiota | GOMC.bin.12837 | g__Cephaloticoccus;s__ | 98.63 | 0.68 | 45 | 14.34 |
| Verrucomicrobiota | GCA_009905215.1 | g__UBA5540;s__UBA5540 sp009905215 | 93.41 | 3.41 | 48 | 13.85 |
| Verrucomicrobiota | GCA_010681825.1 | g__JAAGVW01;s__JAAGVW01 sp010681825 | 99.96 | 0 | 40 | 12.66 |
| Verrucomicrobiota | TARA_ SAMEA2620980_ METAG_ GLODDDOJ | g__Cephaloticoccus;s__Cephaloticoccus sp002713695 | 99.32 | 0 | 34 | 12.32 |
| Verrucomicrobiota | TARA_ SAMEA2621990_ METAG_ NPMDPEGK | g__UBA7441;s__UBA7441 sp002862945 | 98.65 | 0 | 23 | 12.27 |
